# Postnatal xanthine metabolism regulates cardiac regeneration in mammals

**DOI:** 10.1101/2024.07.24.605040

**Authors:** Yuichi Saito, Yuki Sugiura, Akane Sakaguchi, Tai Sada, Chihiro Nishiyama, Rae Maeda, Mari Kaneko, Hiroshi Kiyonari, Wataru Kimura

## Abstract

Postnatal cardiomyocyte cell cycle withdrawal is a critical step wherein the mammalian heart loses regenerative potential after birth. Here, we conducted interspecies multi-omic comparisons between the mouse heart and that of the opossum, which have different postnatal time-windows for cardiomyocyte cell cycle withdrawal. Xanthine metabolism was activated in both postnatal hearts in parallel with cardiomyocyte cell cycle arrest. The pentose phosphate pathway (PPP) which produces NADPH was found to decrease simultaneously. Postnatal myocardial tissues became oxidized accordingly, and administration of antioxidants to neonatal mice altered the PPP and suppressed the postnatal activation of cardiac xanthine metabolism. These results suggest a redox-dependent postnatal switch from purine synthesis to degradation in the heart. Inhibition of xanthine metabolism in the postnatal heart extended postnatal duration of cardiomyocyte proliferation and maintained postnatal heart regeneration potential in mice. These findings highlight a novel role of xanthine metabolism as a metabolic regulator of cardiac regeneration potential.

## Introduction

The mammalian heart lacks the capacity to regenerate itself after injury, with an exception of fetal and early neonatal periods during which a large proportion of cardiomyocytes are proliferative. The vast majority of mammalian cardiomyocytes exit the cell cycle postnatally, which is known to play a central role in the diminishment of the capacity for cardiac regeneration^1^. This is in parallel with multifaceted changes in the myocardium including metabolism, electrophysiology, cellular morphology, among others^2^. Molecular basis of interplays between these physiological transitions and cardiomyocyte cell cycle withdrawal in the postnatal heart has gained an intense research interest. So far, several aspects of environmental and functional transitions at birth have been reported to regulate cardiomyocyte cell cycle exit: namely, exposure to oxygen-rich atmosphere^3^, endothermy acquisition^4^, an increased heart rate^5,6^, and fatty acid intake from milk^7,8^. Interestingly, all of these physiological changes are closely associated with an enhancement of mitochondrial oxidative phosphorylation. Postnatal oxygen-rich environment^3,9^, endothermy acquisition^4^, and faster heart rate^6^ directly or indirectly increase mitochondrial fatty acid utilization, not only enabling efficient ATP synthesis but also resulting in an increase in reactive oxygen species (ROS) production from mitochondria. Recent studies identified that mitochondrial ROS generated through the electron transport chain trigger postnatal cardiomyocyte cell cycle arrest^3,7^, indicating that postnatal metabolic changes are a direct inducer of cardiomyocyte cell cycle arrest. On the other hand, the role of cellular ROS originated from sources other than mitochondria in cardiomyocyte cell cycle regulation remains to be elusive.

During perinatal period, the rate of DNA synthesis in cardiomyocytes fluctuates drastically^10^, implying concurrent substantial alterations in nucleic acid metabolism. Metabolic fluxes of nucleic acids are regulated by a balance between de novo synthesis, salvage, and degradation pathways of purine nucleotides (mainly adenine and guanine nucleotides) and pyrimidine nucleotides (mainly cytosine, thymine and uracil nucleotides). The de novo purine synthesis pathway depends primarily on the pentose phosphate pathway (PPP), whereas de novo synthesis of pyrimidine nucleotides is a combination of the glutamine metabolic pathway and phosphoribosyl diphosphate (PRPP) produced from PPP^11,12^. Purine and pyrimidine nucleotides are recycled through their respective salvage pathways, which also serve as an important source of cellular nucleotides^11,12^. Conversely, excess purine nucleotides are converted into uric acid via hypoxanthine and xanthine, and excess pyrimidine nucleotides are metabolized to acetyl-CoA, for degradation^12^. Xanthine oxidase (XO), a key rate-limiting enzyme that catalyzes the conversion of hypoxanthine to xanthine and xanthine to uric acid through oxidative hydroxylation^13^ serves as a major source of cytosolic ROS production in the context of ischemia-reperfusion and inflammation^14,15^. However, physiological significance of nucleotide metabolism and XO in cardiomyocytes, especially in postnatal cell cycle regulation, has been largely overlooked.

Here, we performed interspecies comparison of the metabolome and transcriptome of the postnatal mouse heart and opossum heart, and found that the purine nucleotide degradation pathway (xanthine metabolic pathway) is commonly upregulated postnatally. Detailed metabolome analysis provided evidences of redox dependent modification of the PPP flux that activates degradation of excess purine nucleotides through the xanthine metabolic pathway. Importantly, inhibition of xanthine oxidase activity reduced oxidative DNA damages and enhanced proliferation in postnatal cardiomyocytes in mice. Moreover, we found that the inhibition of xanthine oxidase activity can extend postnatal regenerative period after myocardial infarction in mice. Our results highlight a novel regulatory link between nucleotide metabolism and redox regulation in the postnatal heart, underscoring the role of cytoplasmic ROS generated through purine nucleotide degradation in the regulation of cardiomyocyte proliferation and cardiac regenerative capacity.

## Results

### Interspecies comparison of the postnatal heart metabolome during cell cycle arrest

To identify unknown metabolic mechanisms regulating postnatal cardiomyocyte cell cycle arrest, we sought to profile the metabolome of postnatal mammalian heart. A couple of studies have conducted comprehensive metabolomic profiling of the postnatal mammalian heart^16,17^. However, identification of metabolic pathways that directly govern cardiomyocyte cell cycle is challenging given the drastic and multifaceted nature of postnatal metabolic switching in the heart. Recently, we found that in a marsupial, the gray short-tailed opossum (*Monodelphis domestica*, hereinafter referred simply to as opossum), cell cycle arrest and loss of regenerative capacity in cardiomyocytes takes place 2-4 weeks after birth, significantly later than that of all other mammals investigated so far^18^. By utilizing this exceptionally long proliferative duration in the opossum heart, we sought to distinguish metabolic pathways that directly regulate cardiomyocyte cell cycle arrest from broader spectrum of postnatal metabolic changes. Our approach is based upon the hypothesis that metabolic pathways that regulate the cell cycle should remain unchanged upon birth, and simultaneously change with cardiomyocyte cell cycle withdrawal in the opossum heart.

We therefore compared metabolomic profiles of the postnatal mouse heart at postnatal day 1 (P1) and P14, and those of the opossum heart at P14 and P28, corresponding to the timings of cardiomyocyte cell cycle arrest in each species (Figure 1A, Supplementary Figure 1A-H, Supplementary Table 1). PCA analysis showed metabolites in opossum hearts have undergone similar types of changes in a postnatal development in mouse hearts (Figure 1B, PC1). Consistent with the changes of cardiomyocyte proliferation rates^18^, in a postnatal development axis (PC1), P14 opossum hearts resembled P1 mouse hearts and P56 opossum hearts resembled P14 mouse hearts (Figure 1B). Almost half of significantly increased metabolites in the mouse heart (Figure 1C) were also increased in the opossum heart (Figure 1D), and vice versa. In contrast, only a few decreased metabolites were shared in both species (Figure 1C-D). The pathway analysis of the commonly altered metabolites revealed a significant change in purine metabolism (Figure 1E).

**Figure 1.**
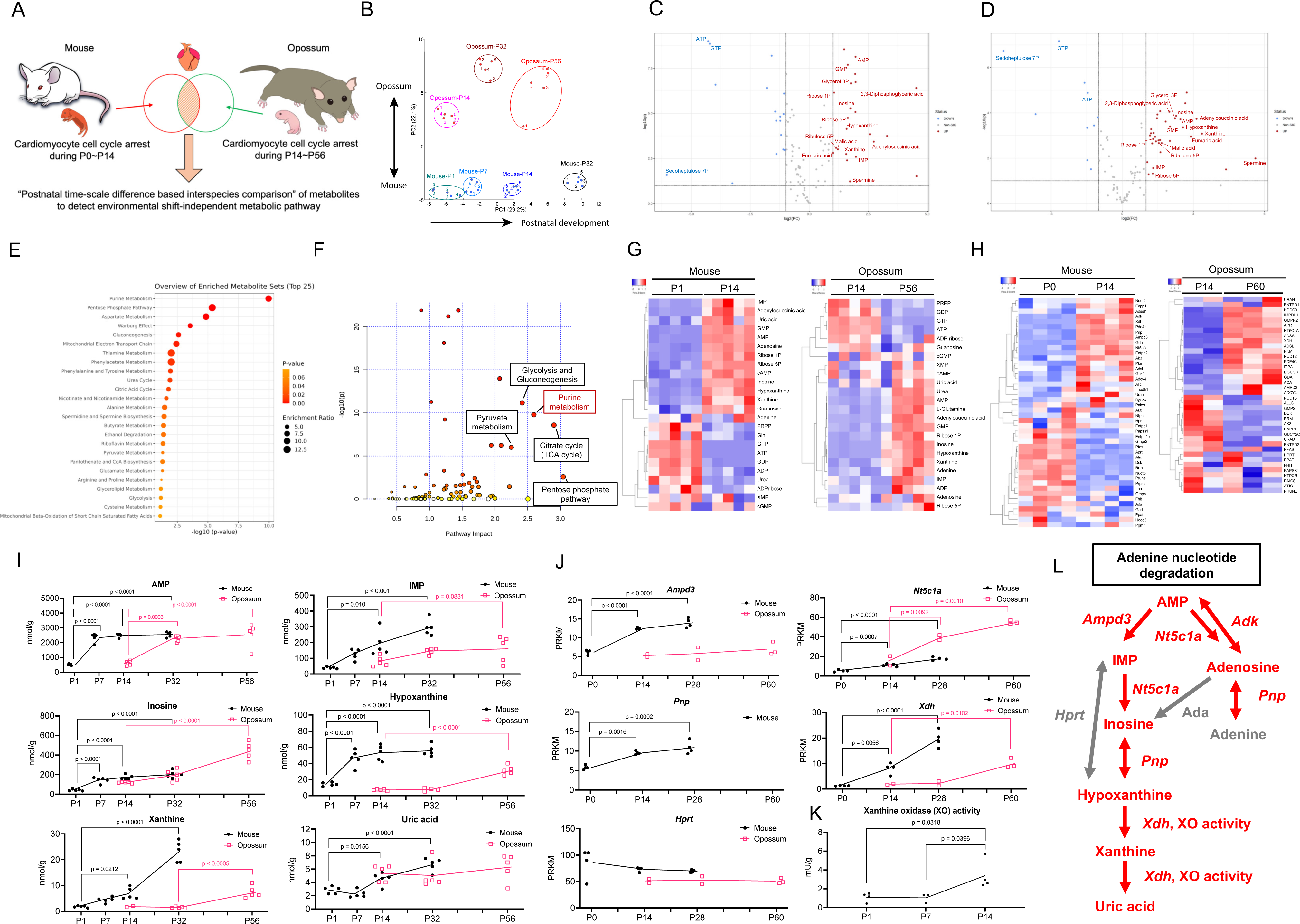
Metabolomic and transcriptomic analyses of the postnatal mouse and opossum heart. (A) A scheme depicting our strategy to identify metabolic pathway(s) that are evolutionary conserved, independent of postnatal environmental changes, and temporally correlated with cardiomyocyte cell cycle arrest. (B) Principal component analysis (PCA) of the comparative metabolome data. (C) Volcano plot of the increased and decreased metabolites in the P14 mouse heart compared with those in P1 mouse heart. Labeled metabolites are commonly increased/decreased in the postnatal opossum heart. (D) Volcano plot of the increased and decreased metabolites in the P56 opossum heart compared with those in the P14 opossum heart. Labeled metabolites are commonly increased/decreased in the postnatal mouse heart. (E) Enrichment pathway analysis of the commonly altered metabolites in the postnatal mouse and opossum heart. (F) Integrated analysis of transcriptome (P0 vs P14) and metabolome data (P1 vs P14) corrected from the postnatal mouse heart. Transcriptome data were reported in Cardoso-Moreira, M. et al^56^. (G) Heatmap representation of the purine metabolism-related metabolites in the postnatal mouse (left) and opossum (right) heart. (H) Heatmap representation of the expression of purine metabolism-related genes in the postnatal mouse (left) and opossum (right) heart. (I) Time-course quantification of purine metabolism-related metabolites. (J) Time-course quantification of purine metabolism-related gene expression (re-analyzed from Cardoso-Moreira, M. et al^56^). (K) Time-course analysis of xanthine oxidase activity in the postnatal mouse heart. (L) A schematic representation of the increased/decreased metabolites and genes related to the purine metabolic pathway from adenine nucleotides to uric acid. Metabolites and genes colored in red are increased/upregulated, and metabolites and genes colored in gray showed no significant changes from P1 to P14 in the mouse heart.

Given metabolic fluxes were not simply reflected in the amount of each metabolite, we conducted integrated-omics analyses of transcriptome and metabolome in the postnatal mouse heart. Top 5 pathways based on the pathway impact scores in the integrated-omics analysis included the purine metabolism, as well as previously reported metabolic pathways associated with cardiomyocyte cell cycle arrest, such as the TCA cycle, glycolysis and pyruvate metabolism^7^ (Figure 1F, Supplementary Table 2). These results prompted us to investigate the dynamics of the purine nucleotide metabolism in the postnatal heart. We firstly compared metabolites (Figure 1G) and genes encoding enzymes involved in the purine metabolism (Figure 1H, some genes lack opossum counterparts) in the postnatal mouse and opossum heart, and demonstrated significant changes in the purine metabolism in both species. Time course quantification of these metabolites, as well as the expression of genes encoding purine metabolic pathway enzymes, revealed that the adenine nucleotide degradation pathway (from adenine to uric acid) were constantly up-regulated postnatally (Figure 1I-J). The only exception was the expression of *Hprt*, which encodes an enzyme that regulates the salvage pathway converting hypoxanthine to IMP (Figure 1J, L). On the other hands, there was an increase in adenosine, and no change in the level of adenine in the P14 mouse heart compared with those in the P1 mouse heart, suggestive of unchanged adenine synthesis from AMP (Supplementary Figure 1I). Also, the expression of *Ada*, which encodes an enzyme regulating the synthesis of inosine from adenosine, was unchanged (Supplementary Figure 1J), suggesting that, unlike xanthine production via IMP, production of xanthine via adenosine is not upregulated from P1 to P14 in the mouse heart. Furthermore, we detected a postnatal increase in the enzymatic activity of XO, from P1 to P14 in the mouse heart (Figure 1K). In summary, interspecies comparison of metabolic changes identified drastic and evolutionary-conserved activation of adenine nucleotide degradation via hypoxanthine and xanthine during cardiomyocyte cell cycle arrest in the postnatal mammalian heart (Figure 1L).

### Postnatal metabolic shift in the PPP

Our data indicate that, during the process of cardiomyocyte cell cycle arrest, adenine nucleotide catabolism is activated without significant changes in its salvage pathway. In order to comprehensively assess purine nucleotide homeostasis, we analyzed de novo purine biosynthesis, which is primarily mediated by the PPP, in the postnatal heart. Interestingly, PPP was the second most significantly altered metabolic pathway in the postnatal heart based on the P values of cross-species metabolomic analysis (Figure 1E), and exhibited the highest Pathway Impact score in the integrated transcriptomic and metabolomic analysis of the mouse heart (Figure 1F). We therefore performed time-course quantification of the PPP intermediate metabolites in the postnatal mouse heart (Figure 2). Most PPP metabolites except for fructose 6-phosphate (F6P) were decreased at P14 compared with those at P1 (Figure 2A-B). This is consistent with the reduced demand for DNA synthesis in the heart as cell proliferation decreases from P1 to P14. F6P is an intermediate metabolite in the pathway that returns to the glycolysis rather than de novo nucleotide synthesis, as shown in a schematic in Figure 2G, suggesting that de novo nucleotide synthesis is decreased in the postnatal heart. The enzyme activity of glucose-6-phosphate dehydrogenase (G6PD), the rate limiting enzyme mediating the initial step of the PPP^19^, was decreased at the P14 mouse heart, providing further support for a decrease in PPP flux (Figure 2C).

**Figure 2.**
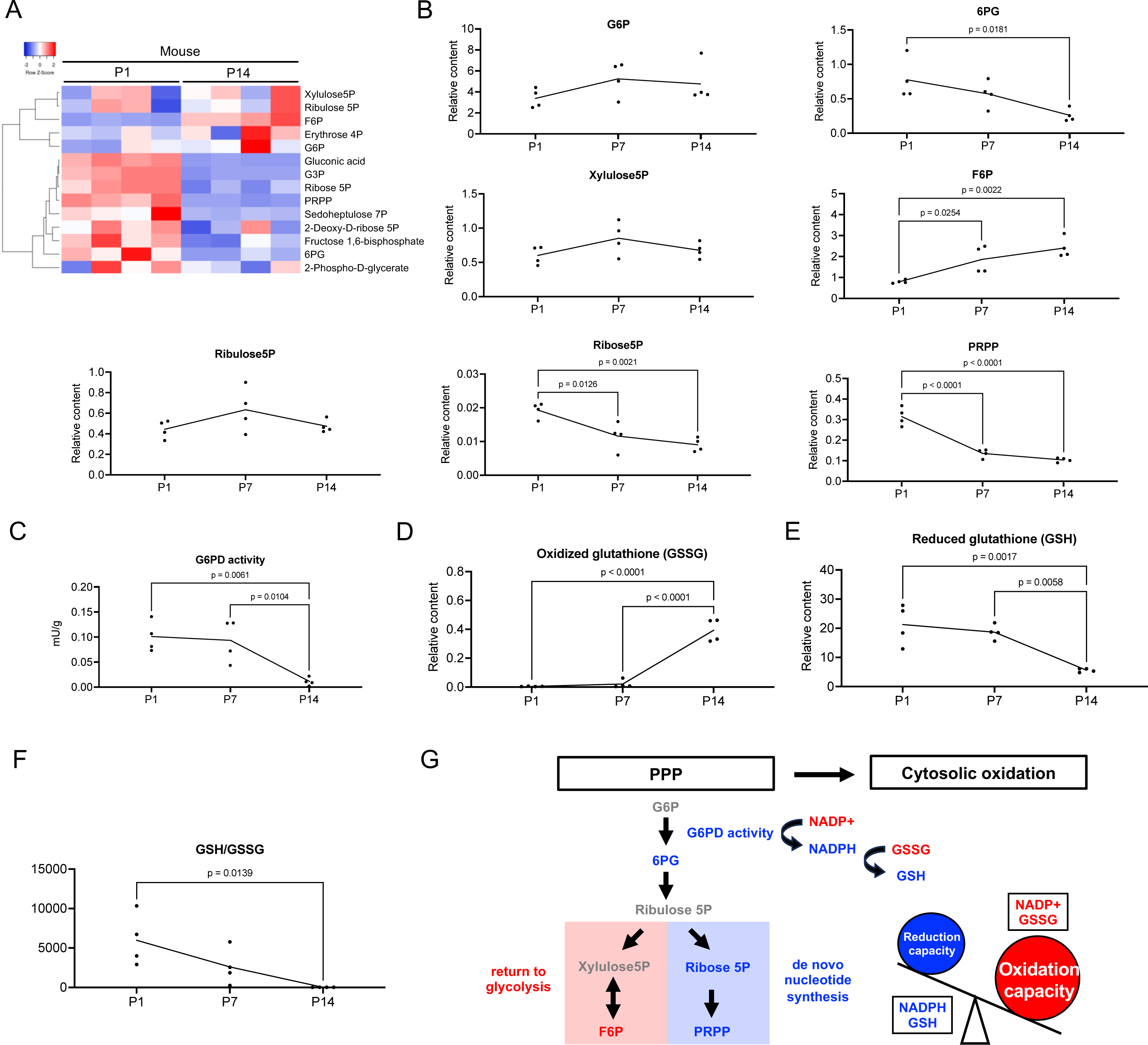
Postnatal alteration of the pentose phosphate pathway in the mouse heart. (A) Heatmap representation of the pentose phosphate pathway (PPP)-related metabolites in the P1 and P14 mouse heart. (B) Time-course quantification of PPP-related metabolites in the postnatal mouse heart. (C) Time-course analysis of glucose-6-phosphate dehydrogenase (G6PD) activity in the postnatal mouse heart. (D-F) Mass spectrometry analysis of (D) oxidized glutathione (GSSG), (E) reduced glutathione (GSH), and (F) ratio of the GSH and GSSG in the postnatal mouse heart. (G) A schematic representation of the postnatal changes in the PPP, and branching metabolic pathways including the generation of NADPH, de novo nucleotide biosynthesis, and returning back to glycolysis. Metabolites colored in red were increased, and those colored in blue were decreased, and those colored in gray showed no significant change in the P14 heart compared with P1.

G6PD not only initiates the first step of de novo purine nucleotide biosynthesis, but also synthesizes NADPH, an essential molecule for the maintenance of the cellular redox state by providing a large part of the cellular reduction potential^20^. We thus hypothesized that redox state in cardiomyocytes shifts towards oxidation after birth due to the reduction in G6PD activity. To test this, we measured the redox state of glutathione, a key physiological antioxidant and indicator of cytoplasmic redox state because the majority of glutathione molecules localize in the cytoplasm^21^. Oxidized glutathione (GSSG) levels were significantly increased, and reduced glutathione (GSH) levels were significantly decreased in the heart over 2 weeks postnatally (Figure 2D-F), as shown in a previous study^3^. These results indicate that the cytoplasmic redox state in the postnatal heart shifts toward oxidation over time, likely due to a decrease in PPP activity. This postnatal downregulation of the PPP may be the result of a decline in the DNA demand as cardiomyocytes are withdrawn from the cell cycle (Figure 2G).

### Xanthine metabolism is regulated by the cellular redox state in the postnatal heart

Our data clearly show that, although de novo purine nucleotide biosynthesis declines as cardiomyocytes exit the cell cycle shortly after birth, purine nucleotide degradation continuously increases for one month after birth in the heart. This led us to hypothesize that, despite the decline in the PPP in the postnatal heart, de novo synthesis of purine nucleotides continues as a byproduct of G6PD-mediated NADPH synthesis in response to the enhanced oxidization state of the myocardial tissue. To assess whether a reduced redox state further downregulates the PPP in the postnatal heart, we injected neonatal mice with N-acetyl cysteine (NAC), a glutathione precursor reported to reduce oxidative state in the postnatal mouse heart^3^ (Figure 3A). Subsequently, we analyzed the metabolomic profile of the P14 heart (Figure 3B). NAC treatment reduced PPP metabolites, including 6-phosphogluconate (6PG), ribulose-5-phosphate (Ribulose5P), xylulose-5-phosphate (Xylulose5P), and ribose-5-phsphate (Ribose5P) (Figure 3C-D). The activity of G6PD showed a tendency toward reduction, which did not reach statistical significance likely due to considerable variation across the samples (Figure 3E). These results suggest that, while postnatal PPP activity decreases in parallel with the reduction of the DNA demand, de novo nucleotide synthesis through the PPP persists due to the need for NADPH in the maintenance of the cytosolic reduction state (Figure 2G and Figure 3F).

**Figure 3.**
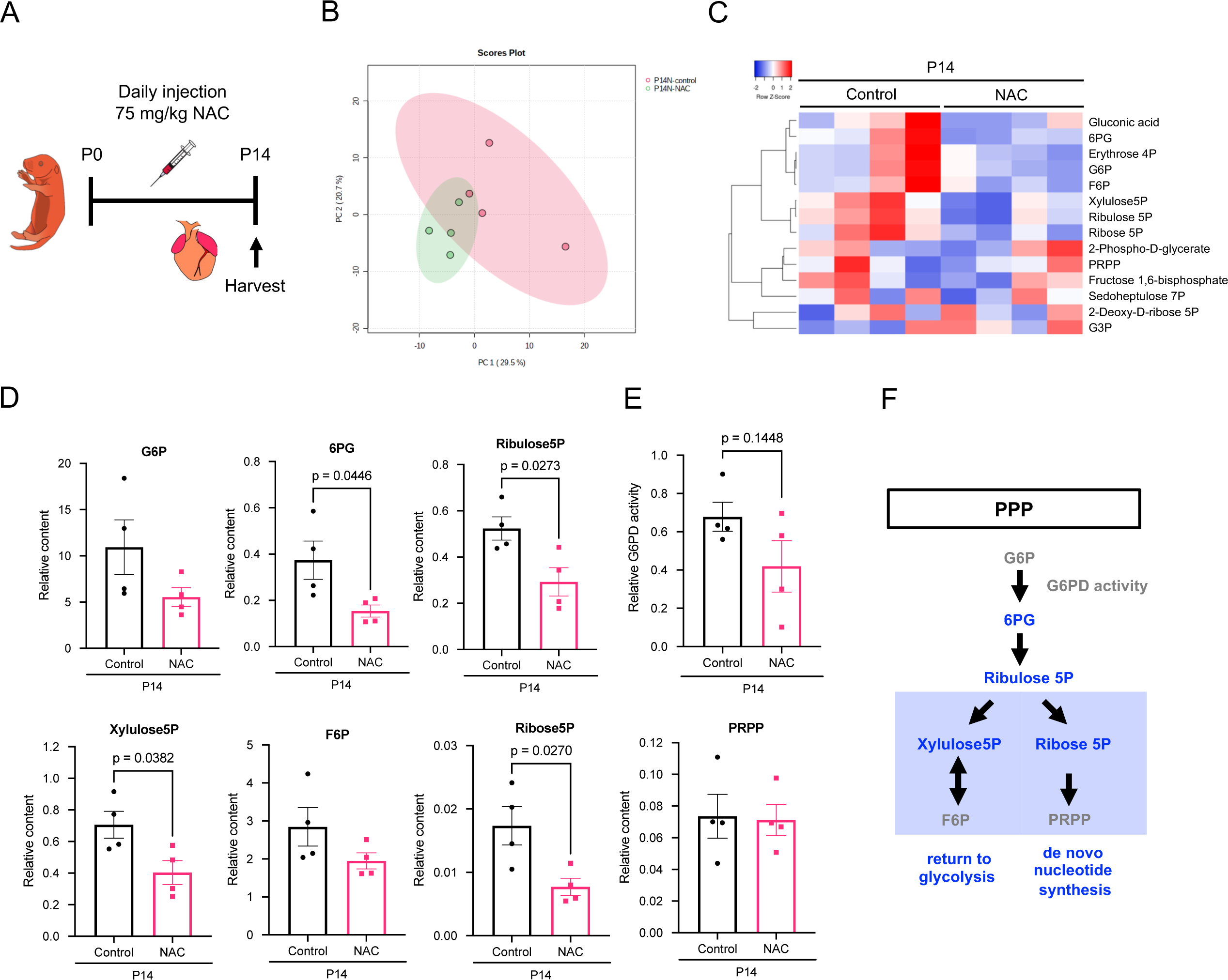
Administration of NAC reduces the PPP in the postnatal mouse heart. (A) A scheme depicting the protocol for a glutathione precursor, N-acetyl cysteine (NAC), injection to neonatal mice. (B) PCA of the metabolomic profiles of the heart harvested from control (red dots) and NAC-treated mice (green dots). (C) Heatmap representation of PPP-related metabolites in the heart of NAC-treated mice at P14. (D) Quantitative data of PPP-related metabolites in the heart of NAC-treated mice. (E) G6PD activity in the heart of NAC-treated mice at P14. (F) A schematic representation of the PPP in the P14 heart from NAC-treated mice. Metabolites colored in blue were decreased, and those colored in gray showed no significant change upon NAC treatment in the P14 heart compared with control.

A previous study showed that cytosolic oxidation is associated with the induction of xanthine metabolism in cold preserved human red blood cells^22^. To test whether oxidation in the postnatal myocardium is a driver of the xanthine metabolism activation in the postnatal heart, we analyzed xanthine metabolism-related metabolites in the NAC-treated P14 heart. NAC treatment significantly altered purine metabolism (Figure 4A). Importantly, some intermediate metabolites of xanthine metabolism, such as IMP or hypoxanthine, were significantly decreased in the NAC-treated heart (Figure 4B). Detailed analysis showed a decrease in most metabolites in the xanthine metabolic pathway in the P14 NAC-treated heart (Figure 4C-D). These results suggest that the reduction of postnatal cardiac redox state with glutathione supplementation reverses postnatal activation of purine nucleotide degradation (Figure 4E), and thus highlight the role of cellular oxidation as a crucial factor driving xanthine metabolism in the postnatal heart.

**Figure 4.**
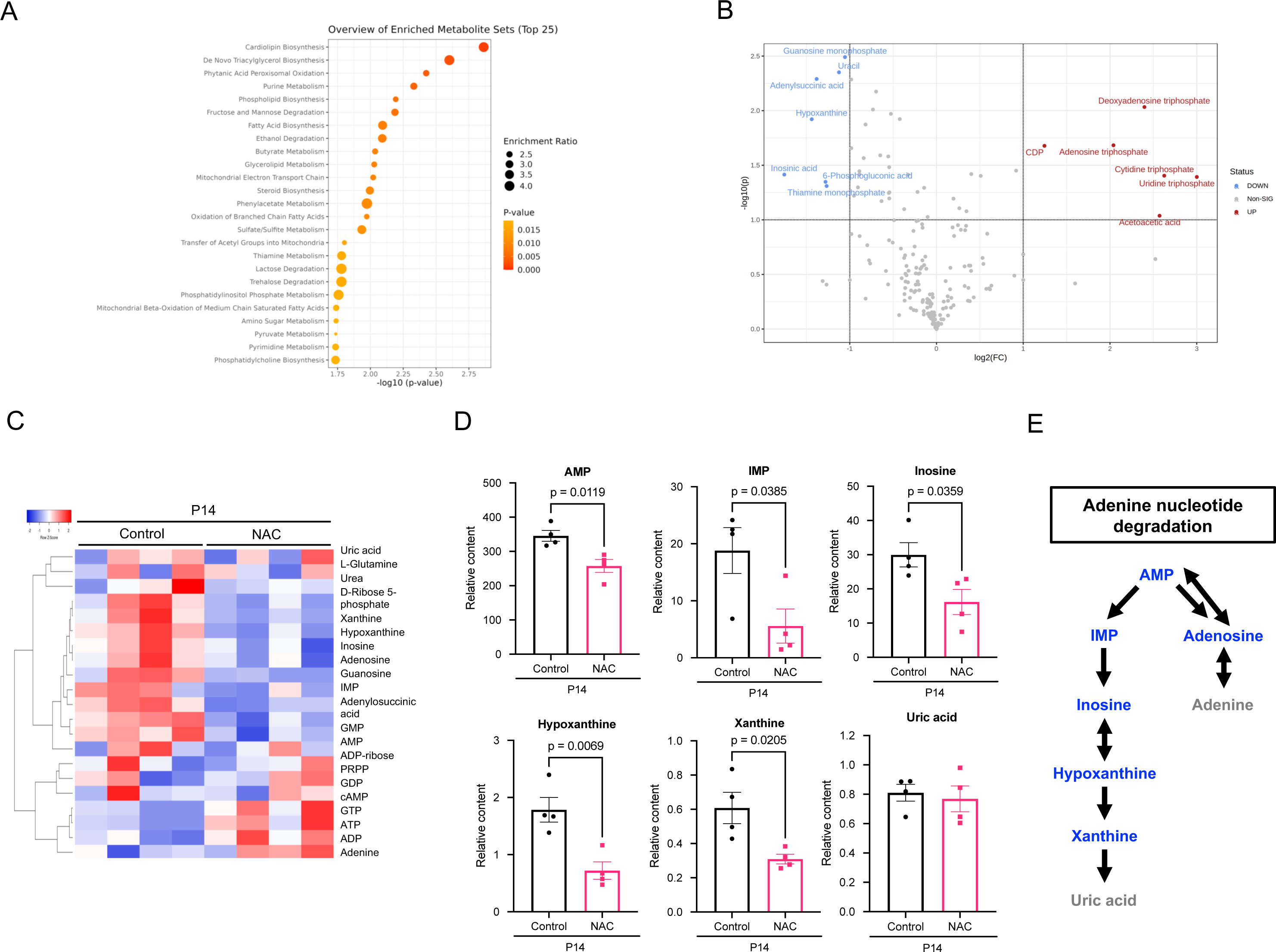
Redox-dependent xanthine synthesis in the postnatal hear. (A) Enrichment pathway analysis of the metabolomic profiles of the heart harvested from control and NAC-treated mice at P14. (B) Volcano plot of the metabolomic profiles of the heart from control and NAC-treated groups. Labeled metabolites were significantly increased (red) or decreased (blue) in the NAC-treated group compared with control. (C) Heatmap representation of the purine metabolism-related metabolites in the heart from control and NAC-treated groups at P14. (D) Quantitative data of purine metabolism-related metabolites in the heart from control and NAC-treated groups at P14. (E) A schematic representation of the adenine nucleotide degradation pathway in the heart from NAC-treated mice. Metabolites colored in blue were decreased, and those colored in gray showed no significant change upon NAC treatment in the P14 heart compared with control.

Mitochondria-derived oxidative stress through fatty acid oxidation is a critical trigger for cardiomyocyte cell cycle arrest in the postnatal heart^7^. We thus assessed the relationships between mitochondrial fatty acid oxidation and xanthine metabolism in the postnatal heart. Pyruvate dehydrogenase kinase isoform 4 (PDK4) plays a central role in the switching from glucose oxidation to fatty acid oxidation by redirecting pyruvates from the cytoplasm to mitochondria in the heart^23,24^. A previous study has demonstrated that a PDK4 inhibitor, dichloroacetate (DCA), enhances glucose oxidation, suppresses mitochondrial fatty acid oxidation, reduces mitochondrial ROS, and induces cell cycle re-entry, in cardiomyocytes of adult mice^7^. This prompted us to investigate whether DCA treatment extends the postnatal duration of cardiomyocyte proliferation in neonatal mice (Supplementary Figure 2A). The number of cardiomyocytes retaining proliferative potential at P14 increased upon DCA treatment (Supplementary Figure 2B-C), suggesting that mitochondrial fatty acid utilization regulates cardiomyocyte proliferation in the neonatal heart. Metabolomic profiling of the DCA-treated heart at P14 showed that DCA mainly affected glucose-alanine cycle^25^, reflecting altered balance of glycolysis and fatty acid oxidation by PDK4 inhibition (Supplementary Figure 3A-D, Supplementary Table 4). However, neither the PPP nor xanthine metabolism was affected by DCA treatment (Supplementary Figure 3E-F). Given that NADPH flux, a critical determinant of cellular reducing power, are regulated independently in the cytosol and mitochondria^26^, activation of the xanthine metabolism in the postnatal heart may be induced by cytoplasmic ROS rather than mitochondrial ROS.

To gain further mechanistic insight into the change of DNA demand and purine metabolism in the postnatal heart, we examined deoxyadenosine triphosphate (dATP), a building block of DNA synthesis^27^ (Supplementary Figure 4). Concurrently with decreased cardiac cell proliferation and DNA demand from P1 to P14, the levels of dATP in the mouse heart were significantly decreased (Supplementary Figure 4A). NAC treatment increased the dATP level in the P14 heart (Supplementary Figure 4B), and DCA treatment also showed a trend towards an increase in the dATP level (Supplementary Figure 4C), in accordance with the extended cardiomyocyte proliferative window^3^ (Supplementary Figure 2). These results indicate that, unlike cellular oxidation state, DNA demand may not determine xanthine metabolism influx (Figure 3, Supplementary Figure 3, Supplementary Figure 4).

We also examined the pyrimidine metabolism in the postnatal mouse heart (Supplementary Figure 5, Supplementary Table 5). Intermediate metabolites of the de novo pyrimidine biosynthesis pathway, such as carbamoyl-aspartic acid, orotic acid, UMP, and TMP decreased from P1 to P14 (Supplementary Figure 5A). Concurrently, the expression of *Tk1*, the gene encoding cytosolic form of thymidine kinase, which mediates the pyrimidine salvage pathway that recycles TMP from thymidine^28^, showed a significant reduction (Supplementary Figure 5B). Thymidine, a substrate for thymidine kinase, also decreased postnatally (Supplementary Figure 5C). In contrast, uridine and uracil, intermediate metabolites in the pyrimidine catabolic pathway increased, while the downstream metabolite dihydrouracil decreased significantly (Supplementary Figure 5D). In summary, pyrimidine metabolism in the postnatal heart shows reduced synthesis and enhanced degradation, aligning with the trend observed in purine metabolism.

### Xanthine oxidase causes cardiomyocyte cell cycle arrest

Previous studies have highlighted the critical role of mitochondrial ROS in the regulation of the cell cycle in postnatal cardiomyocytes^3,7^. Given that xanthine oxidase is a significant source of ROS in cytoplasm^14^, we hypothesized that cytoplasmic ROS generated by xanthine oxidase may induce cell cycle exit in postnatal cardiomyocytes.

Firstly, we assessed the effect of xanthine oxidase inhibition by allopurinol, a purine analog that are widely used in clinical practice^29^, on primary cultured neonatal murine ventricular cardiomyocytes (NMVCs, Supplementary Figure 6A). Allopurinol treatment resulted in a dose-dependent increase in EdU incorporation in cardiomyocytes (Supplementary Figure 6B). Next, we investigated whether the activation of xanthine metabolism contributes to postnatal cardiomyocyte cell cycle arrest *in vivo* (Figure 5A). Uric acid level was decreased by allopurinol treatment in the P14 heart (Figure 5B), suggestive of suppressed xanthine oxidase activity in the heart. Allopurinol treatment did not affect the heart weight at P7 and P14 (Figure 5C, Supplementary Figure 7A), and significantly reduced cardiomyocyte cell size (Figure 5D). Importantly, suppression of xanthine oxidase by allopurinol was sufficient to reduce oxidative stress in postnatal cardiomyocytes (Figure 5E). Furthermore, allopurinol treatment increased cardiomyocyte proliferation at both P7 and P14 (Figure 5F). These results suggest that postnatal activation of xanthine oxidase causes cardiomyocyte cell cycle arrest through an induction of oxidative DNA damage.

**Figure 5.**
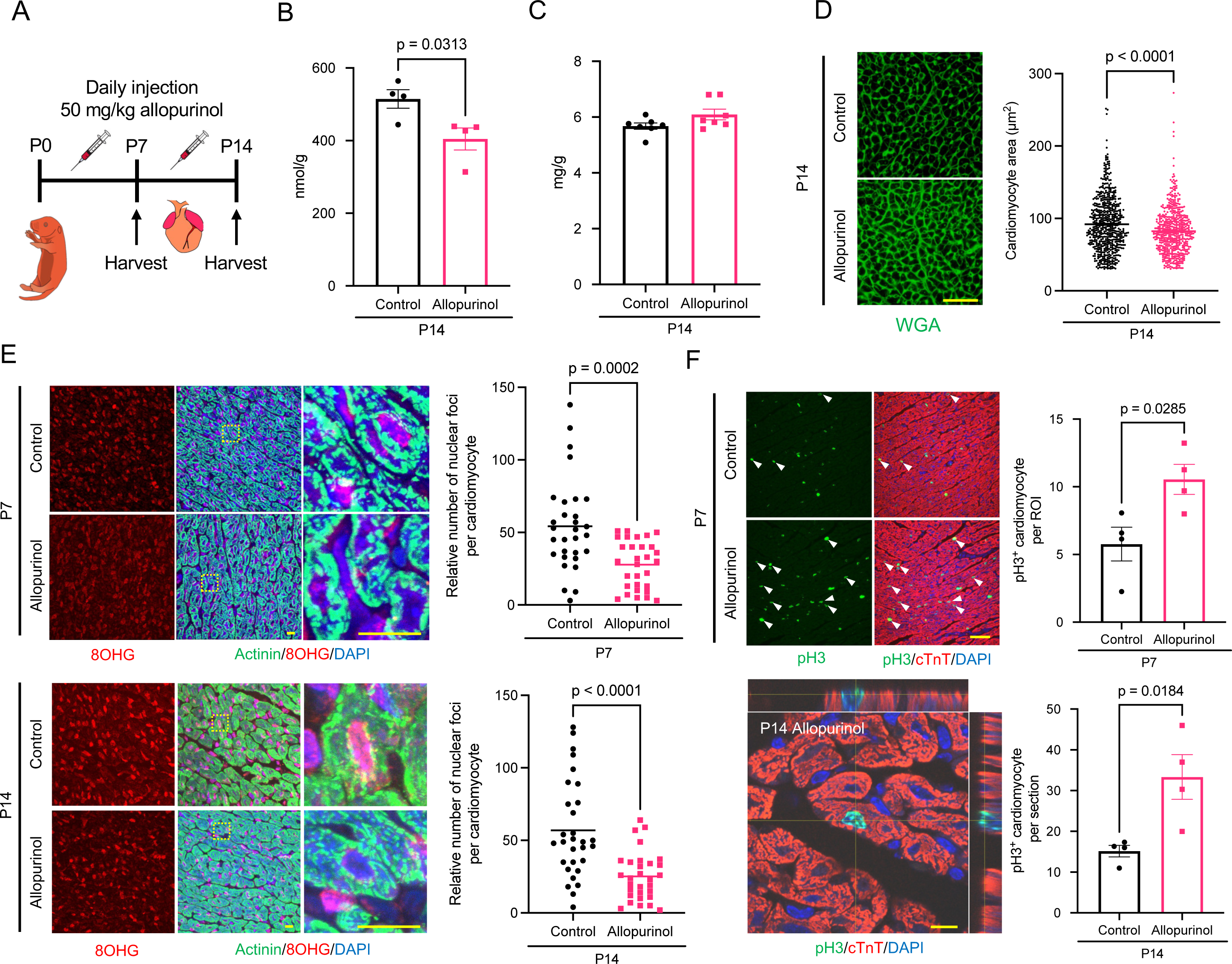
Xanthine oxidase inhibition extends postnatal cardiomyocyte proliferative window. (A) A scheme depicting the protocol of allopurinol administration to neonatal mice. (B) Uric acid levels in the heart from control and allopurinol treated mice at P14. (C) Heart weight per body weight of the P14 heart harvested from control and allopurinol-treated groups. (D) Cardiomyocyte cell size evaluated by the imaging of WGA-stained sections of the heart from control and allopurinol-treated mice. A scale bar indicates 50 µm. (E) Representative images of the immunostaining using antibodies against 8-hydroxyguanosine (8OHG, a marker for oxidative DNA damage), a cardiomyocyte marker α-actinin, and DAPI. Oxidative DNA damage per cardiomyocytes was evaluated by counting the average number of nuclear foci of 8OHG signals in cardiomyocytes. Scale bars indicate 10 µm. (F) Representative images of immunostaining with an anti-phospho-histone H3 (pH3) antibody co-stained with an antibody against cardiac troponin T (cTnT) and DAPI (upper left, white arrowheads indicate pH3+ mitotic cardiomyocytes, a scale bar indicates 50 µm) and confocal z-stack imaging of a pH3+ mitotic cardiomyocyte (bottom left, a scale bar indicates 10 µm). The numbers of mitotic cardiomyocytes per region of interest (ROI, 150 µm x 500 µm) at P7 (top right), and per section at P14 (bottom right) in the LV, septum, and RV were quantified and averaged.

### Inhibition of xanthine oxidase can extend cardiac regeneration window

Given the strong correlation between heart regeneration potential and cardiomyocyte proliferation capacity^1,30–32^, we examined whether the allopurinol treatment maintains regenerative potential in the postnatal heart. We performed daily injection of allopurinol to neonatal mice for 14 consecutive days, and induced myocardial infarction (MI) at P7, by which the regenerative capacity is lost^1^ (Figure 6A). Xanthine oxidase was reported as a critical ROS source that induces myocardial damage following MI and ischemia-reperfusion injury^33,34^. To distinguish acute protective effects of xanthine oxidase inhibition from genuine regenerative response, control mice were also treated with allopurinol for 2 days post-MI (Figure 6A).

**Figure 6.**
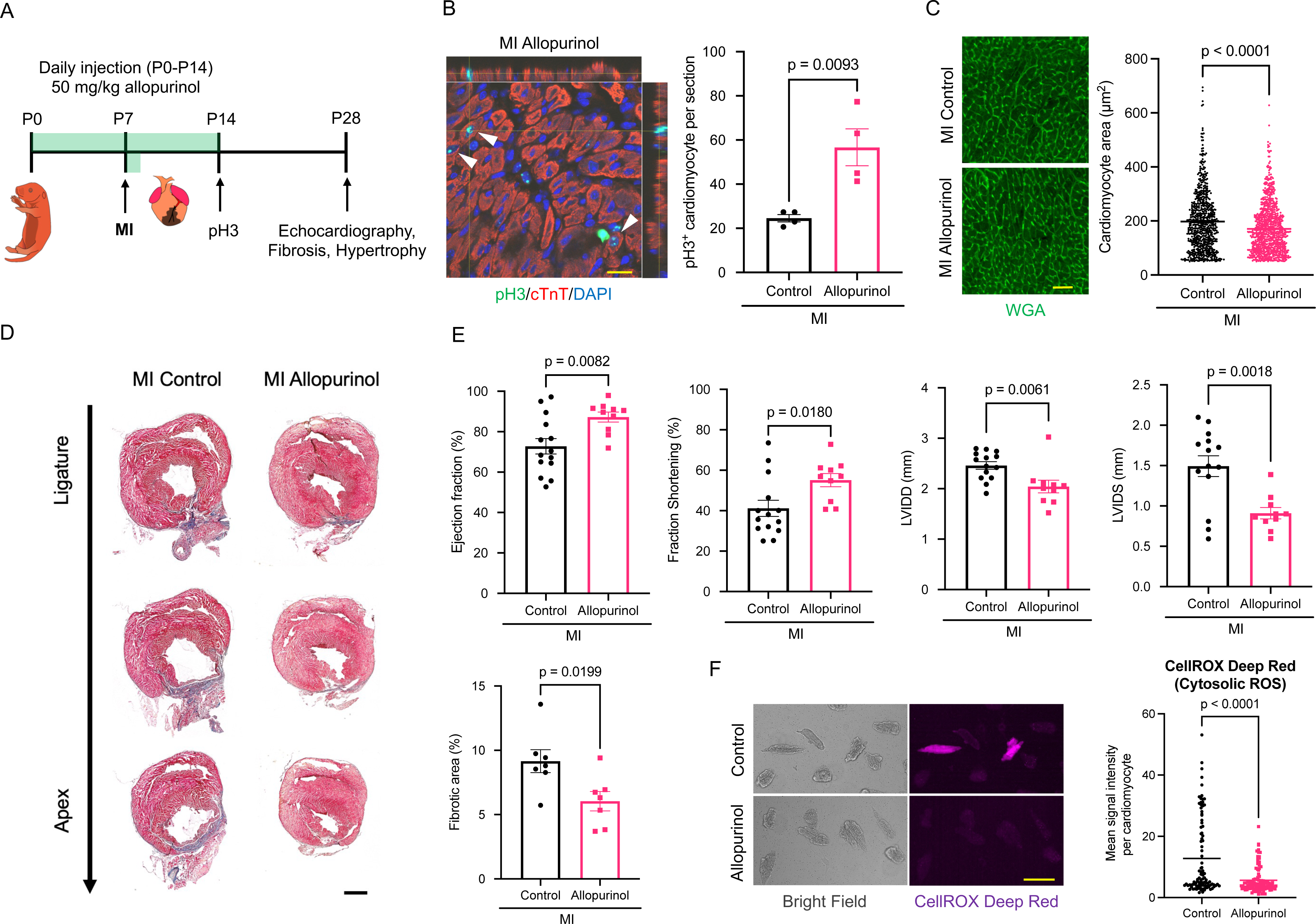
Xanthine oxidase inhibition extends the postnatal window for heart regeneration capacity. (A) A scheme depicting the protocol for daily allopurinol administration, induction of myocardial infarction (MI) to neonatal mice. (B) A representative image of z-stack imaging of the pH3+ cardiomyocytes (left, arrowheads indicate pH3+ mitotic cardiomyocytes) and quantification of cardiomyocyte mitosis at 7 days post injury (dpi). A scale bar indicates 20 µm. (C) Representative images of heart sections stained with WGA (left, a scale bar indicates 50 µm) and quantification of cardiomyocyte cell size at 21 dpi (right). (D) Representative images of Masson’s-Trichrome staining of heart sections and ratio of fibrotic areas and non-fibrotic areas. Analysis was conducted at 21 dpi. A scale bar indicates 1 mm. (E) Echocardiography measurements of ejection fraction (EF), fractional shortening (FS), left ventricular internal diameter diastole (LVIDD), and left ventricular internal diameter systole (LVIDS) at 21 dpi. (F) Representative images of the visualization of cytosolic ROS with CellROX Deep Red after 2-day culture with or without allopurinol and quantification of the cytosolic ROS.

Cardiomyocyte proliferation at 7 days post infarction (dpi) was significantly promoted in the allopurinol-treated group compared with control (Figure 6B). Moreover, at 21 dpi, allopurinol treatment significantly reduced cardiomyocyte hypertrophy (Figure 6C) and cardiac fibrosis (Figure 6D). Consistent with these histological changes, echocardiographic analysis showed a significant improvement in left ventricular systolic function and left ventricular dilatation in the allopurinol-treated group (Figure 6E). To test whether suppression of xanthine oxidase reduces cytosolic ROS level in non-proliferative cardiomyocytes, we isolated and cultured cardiomyocytes from P28 mice and then treated with allopurinol. Allopurinol treated cardiomyocytes showed a significant reduction in cytosolic ROS level (Figure 6F). These results suggest that postnatal xanthine metabolism contributes to the postnatal loss of heart regeneration potential through the production of cytosolic ROS by xanthine oxidase.

## Discussion

Mechanism behind postnatal loss of proliferative and regenerative capacity in cardiomyocytes have gained significant research interest for decades given their potential to enhance myocardial regeneration. The main contributions of this work are three folds: 1) Our multi-omics analysis on the neonatal mouse and opossum heart identified nucleotide metabolism as one of the most significantly altered metabolic pathways during early neonatal period in the mammalian heart. 2) We have demonstrated that nucleotide synthesis and degradation dynamics is coupled with the myocardial redox state in the postnatal heart. 3) Pharmacological intervention has demonstrated that targeting purine nucleotide degradation enhances cardiomyocyte proliferation and extends postnatal duration of cardiac regeneration in neonatal mice. Collectively, our study identified a causative, not merely correlative, link between nucleotide metabolism, redox regulation, and regenerative capacity in the mammalian postnatal heart.

Several previous studies have conducted metabolome analyses and identified alterations in nucleotide metabolism among multitude of changing metabolic pathways in the postnatal mammalian heart^16,17,35,36^. Similarly, our study detected numerous metabolic pathways that significantly alter after birth in the mouse heart (Supplementary Figure 1E). This may reflect the metabolic adaptation to extrauterine environment in the heart, making it challenging to highlight metabolic pathway(s) that causatively link to cell cycle regulation. We therefore took advantage of the delayed postnatal cell cycle arrest in opossum cardiomyocytes^18^, which supposedly enables to discriminate metabolic pathways that causatively link to cardiomyocyte cell cycle arrest, which takes place more than 2 weeks after birth. Our analysis of the metabolites that increased or decreased in both mouse and opossum hearts during cardiomyocyte cell cycle arrest identified nucleotide metabolic pathways as the most significantly enriched metabolite sets (Figure 1E). Our approach thus highlights the utility of exploiting developmental heterochrony between mammalian species to identify novel, evolutionary conserved, and physiologically relevant biological pathways.

Previous studies have suggested the role of the nucleotide biosynthesis pathway on cardiomyocyte cell cycle regulation and cardiac regeneration. The isoenzyme pyruvate kinase muscle isoenzyme 2 (PKM2) has been implicated in regulating cardiomyocyte proliferation and cardiac regeneration through the activation of G6PD and the transcriptional activation of cell cycle-related genes together with β-catenin^37^. Similarly, enhancing glucose uptake in cardiomyocytes has been shown to accelerate cardiac regeneration via nucleotide synthesis in neonates^38^. Cardiomyocyte maturation is also regulated by the PPP in human embryonic stem cell-derived cardiomyocytes (hESC-CMs)^39^. Our study introduces a novel concept where a balance between the demands for NADPH regeneration and nucleotide synthesis is crucial in regulating the cardiomyocyte cell cycle through cytoplasmic ROS production. In the postnatal mammalian heart, the demand for nucleotides decreases as DNA synthesis declines in cardiomyocytes^10^. In line with this, our data indicate a significant suppression of the PPP, a pathway accounting for de novo nucleotide biosynthesis, in the postnatal heart. Since the PPP is the primary cytosolic source of NADPH^40^, downregulation of the PPP may limit NADPH regeneration in cytoplasm. This is also supported by our data showing a postnatal shift in the myocardial redox state towards oxidation. Therefore, the postnatal heart may face conflicting demands for the PPP— reducing nucleotide synthesis while needed to accelerate NADPH regeneration. Our observation indicates an increase in reversion to glycolysis from PPP, which aligns with such contradicting demands. This contradiction might lead to an excess of nucleotide synthesis over demand in the postnatal heart. Our evidence supports this scenario: the purine nucleotide degradation pathway (xanthine metabolic pathway) was constantly upregulated postnatally in the myocardium, indicating that nucleotide synthesis exceeds its requirement. Furthermore, preventing myocardial oxidation by administering an antioxidant NAC reduced PPP activity for both nucleotide biosynthesis and reversion to glycolysis, as well as xanthine metabolic pathway intermediates. This indicates that postnatal oxidation stimulates the need for the PPP and subsequent nucleotide degradation. Despite these changes, NAC administration did not significantly alter NADPH levels (Supplementary Table 1, Supplementary Table 5), possibly due to the independent regulation of mitochondrial and cytoplasmic NADPH pools^26,41^.

Interestingly, our data indicate that the degradation of excess nucleotides further shifts the myocardial redox state towards oxidation via xanthine oxidase after birth, suggesting an imbalance between redox state and nucleotide homeostasis initiates a positive feedback that exacerbates oxidation state in the heart.

ROS are critical mediators in various biological signal transduction processes^42^. Biological roles of ROS vary depending on their sources, and as such, their cellular localization is strictly controlled^42^. It has been known that in the postnatal mouse heart, cardiomyocyte cell cycle withdrawal is regulated by mitochondrial ROS^43,44,45,46^, which induce oxidative DNA damage and activate the DNA damage response pathway^3^. On the other hands, the role of cytosolic ROS on postnatal cardiomyocyte cell cycle arrest has been neglected in previous studies. Our findings indicate that cytosolic ROS derived from xanthine oxidase induce oxidative damage in cardiomyocyte nuclei, resulting in a reduction in cardiomyocyte proliferation and regenerative capacity in the postnatal heart. Xanthine metabolism is well-characterized in the context of ischemia-reperfusion-derived purine nucleotide breakdown^47^, which is generally considered as a pathological ROS source^14^. A previous study showed that allopurinol attenuated left ventricular remodeling and dysfunction after MI in adult mice due to protection of myocardium from oxidative stress^34^. Our study suggests a promising therapeutic potential of allopurinol for inducing myocardial regeneration through cardiomyocyte proliferation. Although several chemicals have been reported to extend the postnatal duration of cardiac regeneration in neonates^3,7,45^ or induce cardiac regeneration in adults^48–50^, allopurinol offers a unique advantage in safety given its long history of widespread use^29^. Yet, further validation and investigation are necessary to assess the safety and efficacy of targeting xanthine oxidase to regenerate the myocardium in mammals.

## Resource availability

### Lead contact

Further information and requests for resources and reagents should be directed to and will be fulfilled by the lead contact: Dr. Wataru Kimura.

### Materials availability

This study did not generate new unique reagents.

## Methods

### Mice

All animal experiments are approved by the Institutional Animal Care and Use Committee (IACUC) of the RIKEN Kobe Branch. All animal experiments were performed on age-matched CD1 (ICR) mice (provided by the Laboratory for Animal Resources and Genetic Engineering in RIKEN and by Oriental Yeast Co., Ltd.) or opossums (provided by the Laboratory for Animal Resources and Genetic Engineering in RIKEN).

### Drug Administration

N-Acetyl-L-cysteine (Sigma, A7250) was reconstituted in PBS. Allopurinol (Wako, 011-12501) and sodium dichloroacetate (DCA, Sigma, 347795) were reconstituted in dimethyl sulfoxide (DMSO) diluted at 1:10 in PBS. NAC, allopurinol, DCA were injected subcutaneously at a daily dose of 75 mg/kg, 50 mg/kg, or 100 mg/kg, respectively, to CD1 mice from postnatal day 0 (P0) to P7 or P14. For the experiment of myocardial infarction (MI), Allopurinol was injected to CD1 mice from P0 to P14, and MI was surgically induced at P7. Allopurinol was injected to control mice at P7 after the MI induction, and also at P8 and P9. Echocardiography was performed at P28 and then the heart was harvested.

### Surgical Induction of Myocardial Infarction in Neonatal Mice

Neonates were placed in a vinyl bag with air holes and anesthetized by hypothermia, induced by placing the bag on ice. MI was induced in neonatal mice at P7 by permanent ligation of the left anterior descending coronary artery with 6-0 polypropylene suture (Ethicon, EP8707) following anesthesia and a lateral thoracotomy. The incision on the thoracic wall was sutured with 6-0 polypropylene, the external incision was sealed with skin adhesive (3M Vetbond), and the neonates were warmed on a thermal plate (42°C) until resuscitation. Left ventricular systolic function was measured by echocardiography using Affiniti50 with an L15-7io transducer (Phillips) performed on non-sedated mice.

### Immunohistochemistry

Tissues were fixed overnight at 4°C with 4% paraformaldehyde (PFA) in PBS and then processed for cryosectioning. Tissues were embedded in tissue-freezing medium and cut to 8 μm thickness. Following antigen retrieval with either 1 mM EDTA/0.05%

Tween 20 in boiling water or epitope retrieval solution (IHC World) with a steamer (IHC-Tek Epitope Retrieval Streamer Set), sections were blocked with 10% serum from the host animal of secondary antibodies, and incubated with primary antibodies overnight at 4°C. Sections were subsequently washed with PBS and incubated with corresponding secondary antibodies for 1 hour at room temperature. For 8OHG staining, sections were blocked with M.O.M. Immunodetection Kit, Basic (Vector Laboratories), and were incubated with primary antibodies for 2 days at 4°C, and then incubated with alpaca recombinant secondary antibodies overnight at 4°C. The primary antibodies used are as follows: Ser10-phosphorylated histone H3 (Millipore, 06–570; 1:100 dilution), cardiac troponin T (BD Pharmingen, 564766; 1:250), anti-sarcomeric α-actinin (Abcam, ab68167; 1:100), anti-8-hydroxyguanosine (Abcam, ab62623; 1:50). Immune complexes were detected with Alexa Fluor 488– or Alexa Fluor 555–conjugated secondary antibodies (Invitrogen), or Alexa Fluor 488– or Alexa Fluor 568–conjugated alpaca recombinant secondary antibody (Invitrogen) or Alexa Fluor 647–conjugated wheat germ agglutinin (ThermoFisher Scientific, W32466; 50 µg/mL) and nuclei were stained with 4′,6-diamidino-2-phenylindole (DAPI) (Nacalai Tesque, 11034-56). The slides were mounted with PLUS antifade mounting medium (Vector Laboratories). Histological images were captured using a BX53 (Evident) or APX100 (Evident) microscope, and confocal images were obtained with an LSM800 microscope (Zeiss).

### Cardiomyocyte cell size quantification

Heart sections were stained with Alexa Fluor 488–conjugated wheat germ agglutinin (ThermoFisher Scientific, W11261; 50 µg/mL) to visualize cell edges and imaged with BX53 or APX100 across the base, apex and middle sections of the left ventricle, right ventricle and septum. Cardiomyocyte size was quantified using ImageJ FIJI software^51^ (National Institutes of Health) by analyzing 500–1000 cells per group.

### Masson’s trichrome staining

Masson’s trichrome staining was performed according to the manufacturer’s protocol (Scy Tek Laboratories, TRM-1) using 4% PFA-fixed cryosections. Histological images were captured using a APX100 (Evident) microscope. Quantification of the ratio of the fibrotic area in trichrome-stained sections was performed using ImageJ FIJI software. In each mouse, the average ratio of the fibrotic area was quantified in three sections from the ligature to the apex at 600 µm intervals.

### Metabolome Analysis

For postnatal interspecies metabolome comparison of the mouse and opossum heart (Table S1), the hearts were harvested and snap frozen. These frozen samples were analyzed by capillary electrophoresis-time of flight mass spectrometry (CE-TOFMS, Basic Scan, Human Metabolome Technologies Japan). Metabolite concentrations were calculated by absolute quantification. Details are described in table S1. For NAC or DCA -treated P14 mouse heart, or postnatal mouse hearts (Supplementary Table 3-5), the heart samples were immediately freeze clamped by pre-chilled freezing clamp (Natsume Seisakusho, KN-838-S). Details are described previously^52^. Briefly, metabolites were detected using an orbitrap-type MS instrument (Q-Exactive focus; Thermo Fisher Scientific) connected to a high-performance IC system (ICS-5000 +, Thermo Fisher Scientific) and a triple-quadrupole mass spectrometer (LCMS-8060, Shimadzu Corporation, Kyoto, Japan). Obtained metabolome data were analyzed using metaboanalyst 5.0 software^53^. Heatmaps were generated by using Heatmapper^54^.

### Enzyme activity assay and uric acid assay

Enzyme activities were evaluated using freeze-clamped mouse heart samples. Xanthine oxidase activity was measured with the Amplex™ Red Xanthine/Xanthine Oxidase Assay Kit (ThermoFisher Scientific, A22182), G6PD activity was measured with the Glucose-6-Phosphate Dehydrogenase Activity Assay Kit (Sigma, MAK015), and uric acid levels in heart lysates were measured with the Uric Acid Assay Kit (Sigma, MAK077), according to the corresponding manufacturers’ protocol. Fluorescence and absorbance were measured by a multimode microplate reader (TACAN, Infinite 200).

### Cardiomyocyte cell culture

Neonatal murine ventricular cardiomyocytes were isolated from P1 mice (10-15 hearts were used per isolation) with the Neonatal Heart Dissociation Kit, mouse and rat (Miltenyi Biotec, 130-098-373), according to the manufacturer’s protocol with some modifications. Briefly, fibroblasts were removed by 2 times of transient seeding on a non-coated dish and incubation for 1.5 hours. Then isolated cardiomyocytes were seeded on fibronectin (Fibronectin Solution human, PromoCell, D13121) coated 96 well plates and cultured for 2 days with allopurinol or control solvent (0.1% DMSO/PBS).

Non-proliferative mouse cardiomyocytes were isolated from P28 mice (1 heart was used per isolation) according to a previously reported protocol^55^. Isolated cardiomyocytes were cultured for 2 days with allopurinol or control solvent (0.1%

### EdU incorporation assay and immunocytochemistry

For EdU incorporation-based primary cardiomyocyte proliferation assay, Click-it EdU Imaging Kit (ThermoFisher Scientific, C10337) was used according to the manufacturer’s protocol. Briefly, cultured neonatal cardiomyocytes were incubated with the culture medium containing 10 µM EdU solution for 2 days, and then cardiomyocytes were fixed by 4% PFA for 10 min. After fixation, cells were blocked with 5% serum from the host animal of the secondary antibody, and incubated with anti-Pericentriolar material 1 (Sigma, HPA023370; 1:2000 dilution) primary antibody overnight at 4°C. Cells were subsequently washed with PBS and incubated with Alexa Fluor 488–conjugated secondary antibody (Invitrogen) for 1 hour at room temperature. After immunocytochemistry, EdU labeling was applied according to the manufacturer’s protocol, and nuclei were stained with DAPI. Cell images were captured using an APX100 (Evident) microscope. The number of cardiomyocyte nuclei surrounded by PCM1 was quantified using ImageJ FIJI software^51^ (National Institutes of Health). Subsequently, the number of EdU^+^ cardiomyocyte nuclei within a defined area was counted to calculate frequency of EdU^+^ cardiomyocytes.

### Cytosolic ROS assay

Cytosolic ROS was assessed by using the CellROX Deep Red reagent (ThermoFisher Scientific, C10422) according to the manufacturer’s protocol. Briefly, cultured juvenile cardiomyocytes were incubated with the culture medium containing 5 µM of CellROX Deep Red reagent for 30 min. After CellROX staining, the medium was replaced with FluoroBrite™ DMEM (Gibco, A1896701) and the images were captured by APX100 (Evident). Mean signal intensity of CellROX Deep Red was measured using ImageJ FIJI software^51^ (National Institutes of Health). Fluorescent signals in rod-shaped cardiomyocytes were quantified across multiple wells for each group.

### Quantifications and statistical analysis

Unless the figure legend indicates otherwise, statistical assessments were carried out with GraphPad Prism 9, applying unpaired t-test for comparison between two groups and one-way analysis of variance (ANOVA) with Tukey’s test for comparison among multiple groups. Regarding comparisons across different species, statistical analysis was as the mean ± SEM. There were no exclusions of data or samples throughout the study.

## Data and code availability

This paper does not report original code.

## Acknowledgments

The authors thank the RIKEN animal facility for animal husbandry. This work was supported by grants from Japan Society for the Promotion of Science (KAKENHI 20K22751 and 21K15356), the Takeda Science Foundation, RIKEN BDR (Center for Biosystems Dynamics Research)-Otsuka Pharmaceutical Co., Ltd. Kakehashi/Bridge program, RIKEN Incentive Research Projects to Y. Saito; grants from the Japan Agency for Medical Research and Development (23zf0127007s0102, JP23zf0127003 and JP23gm1210009) to Y. Sugiura; and grants from Japan Society for the Promotion of Science (KAKENHI 17H05083, 19K22629, 20H03680, and 21K18273), the PRIME program of the Japan Agency for Medical Research and Development, Takeda Science Foundation, Uehara Memorial Foundation, Japanese Circulation Society, Senri Life Science Foundation, Princess Takamatsu Cancer Research Fund, Mitsubishi Foundation, Bristol-Myers Squibb, Mochida Memorial Foundation, Naito Foundation, Toray Science Foundation, and Hoansha foundation, and a RIKEN CDB/BDR intramural grant to W.K.

## Author contributions

Conceptualization, Y. Saito and W.K.; methodology, Y. Saito, Y. Sugiura, A.S., T.S., C.N., R.M., M.K., H.K. and W.K.; formal analysis, Y. Saito, Y. Sugiura, A.S., and W.K..; investigation, Y. Saito, Y. Sugiura, A.S., T.S., C.N., R.M. and W.K.; resources, M.K. and H.K.; writing – original draft, Y. Saito; writing – review & editing, Y. Sugiura, A.S. and W.K.; visualization, Y. Saito, Y. Sugiura, A.S., W.K; supervision, H.K. and W.K.

## Supplementary Figure Legends

**Supplementary Figure 1.**
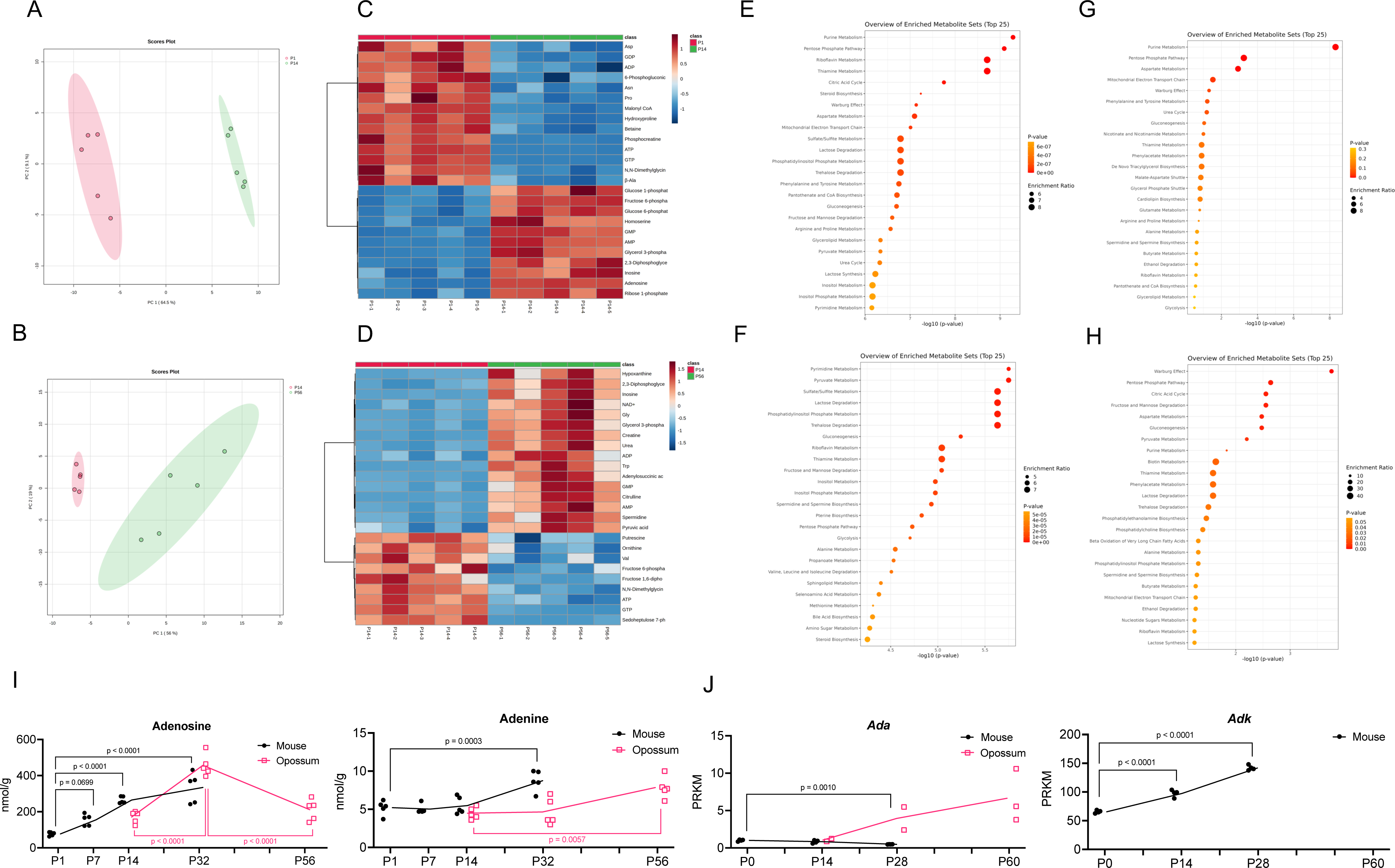
Metabolome analyses of the neonatal mouse and opossum heart during cell cycle arres. (A) PCA of P1 and P14 mouse heart metabolome. (B) PCA analysis of P14 and P56 opossum heart metabolome. (C) Heatmap of the top 25 most significantly increased or decreased metabolites in the mouse heart at P14 compared with P1. (D) Heatmap of the top 25 most significantly increased or decreased metabolites in the opossum heart at P56 compared with P14. (E) Enrichment pathway analysis of significantly increased and decreased metabolites in the mouse heart at P14 compared with P1. (F) Enrichment pathway analysis of significantly increased and decreased metabolites in the opossum heart at P56 compared with P14. (G) Enrichment pathway analysis of increased metabolites both in the mouse and opossum heart postnatally. (H) Enrichment pathway analysis of decreased metabolites both in the mouse and opossum heart postnatally. (I) Time-course quantification of adenosine to adenine nucleotide metabolism-related metabolites. (J) Time-course quantification of adenosine to adenine nucleotide metabolism-related gene expression (re-analyzed from Cardoso-Moreira, M. et al^56^).

**Supplementary Figure 2.**
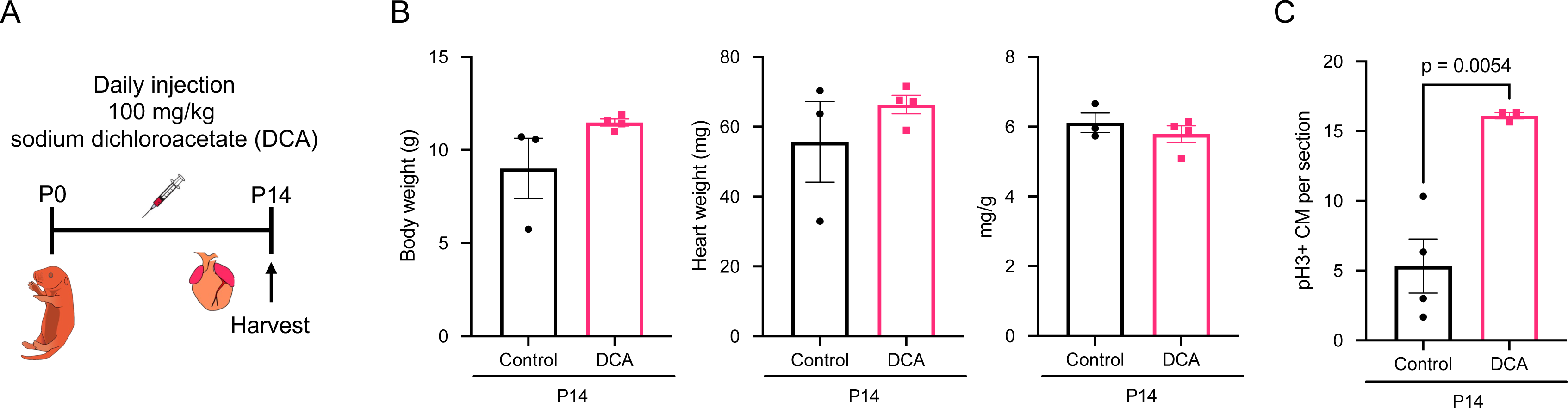
Inhibition of postnatal fatty acid metabolism extended postnatal cardiomyocyte cell cycle window. (A) A scheme depicting the protocol of dichloroacetate (DCA) administration to neonatal mice (B) Body weight, heart weight, and the ratio of heart weight per body weight at P14 in control and DCA-treated mice. (C) Quantification of pH3+ mitotic cardiomyocytes at P14 in control and DCA-treated mice.

**Supplementary Figure 3.**
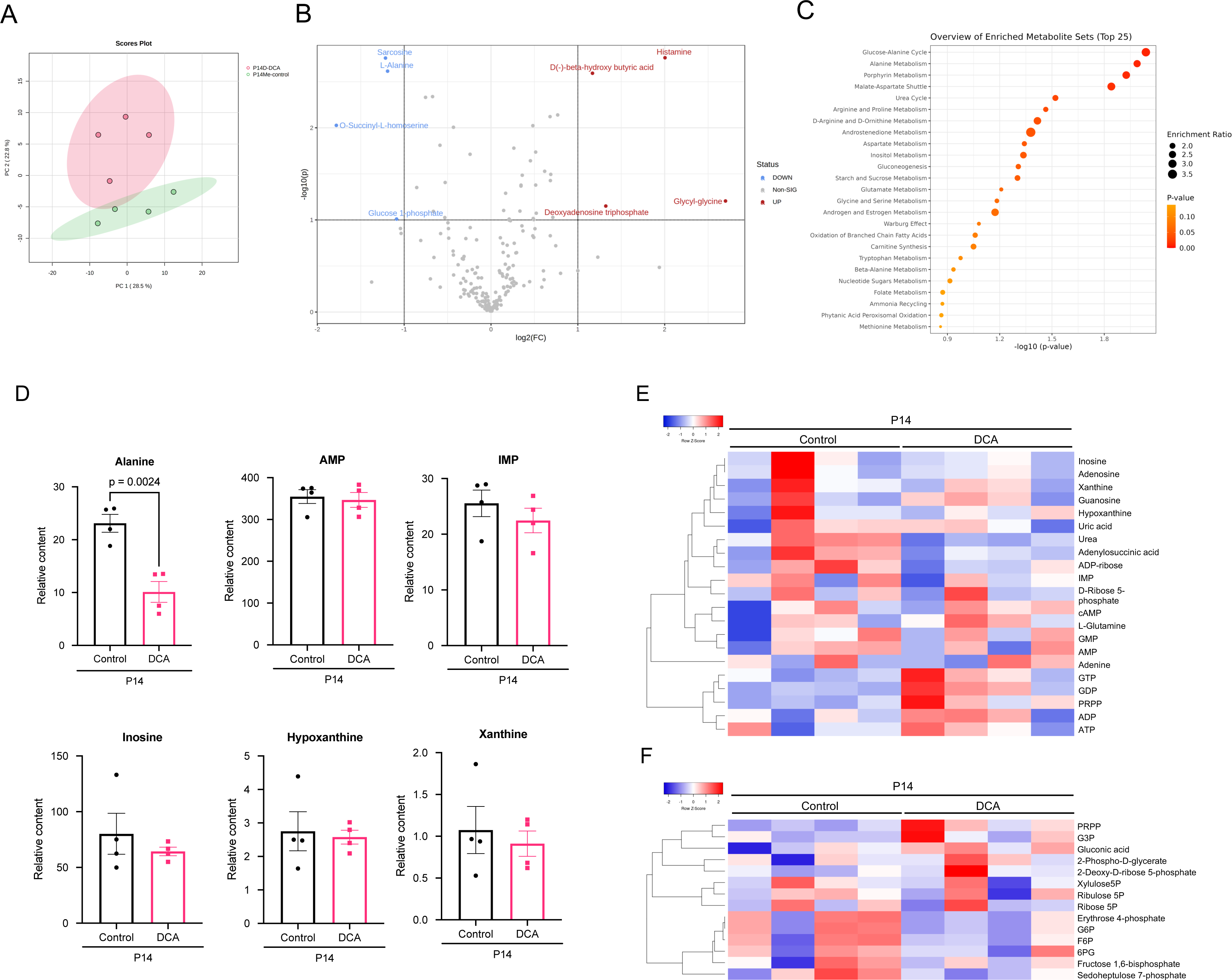
Inhibition of postnatal fatty acid oxidation did not alter xanthine metabolism. (A) PCA of the metabolome data of the heart harvested from control and DCA treated mice at P14. (B) Volcano plot of the metabolites in the heart from DCA treated mice at P14. Significantly increased or decreased metabolites are labeled. (C) Enrichment pathway analysis of significantly increased or decreased metabolites in the heart from DCA treated mouse at P14. (D) Quantitative data of alanine levels (top left) and purine metabolism-related metabolites in the heart from DCA-treated mice at P14. (E) Heatmap of purine metabolism-related metabolites in the heart from DCA-treated mice at P14. (F) Heatmap of PPP-related metabolites in DCA treated mice at P14.

**Supplementary Figure 4.**
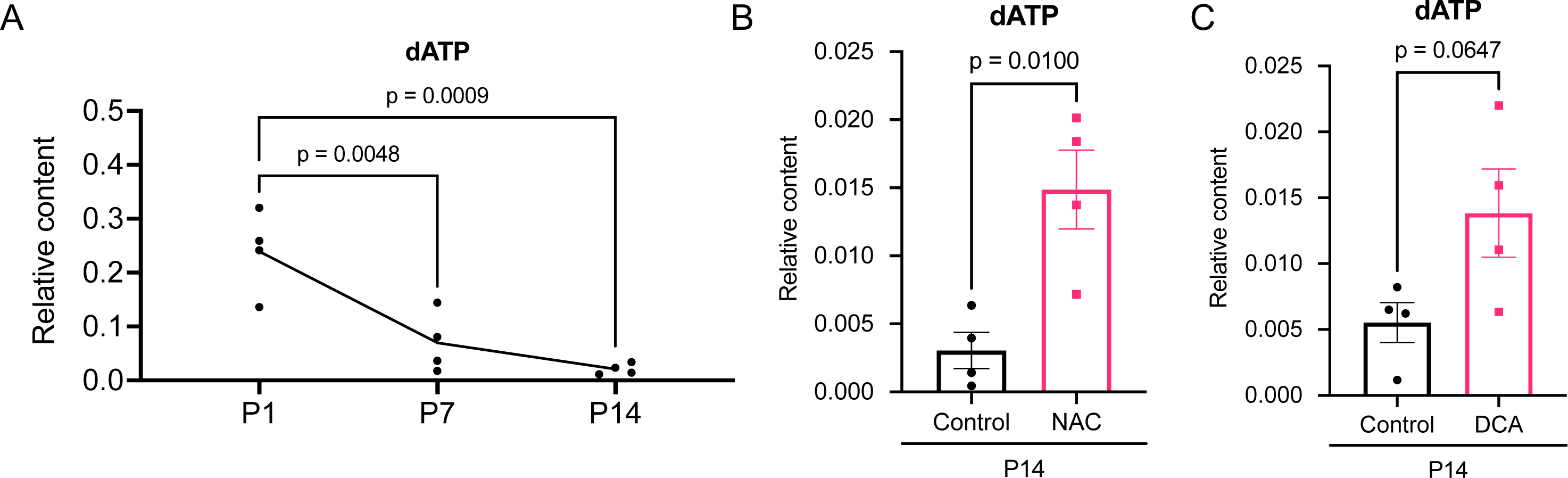
Postnatal changes of adenine-derived DNA building block. (A) Time-course analysis of dATP relative contents in the postnatal mouse heart from P1 to P14. (B) Relative contents of dATP in the heart from NAC-treated mice at P14. (C) Relative contents of dATP in the heart from DCA-treated mice at P14.

**Supplementary Figure 5.**
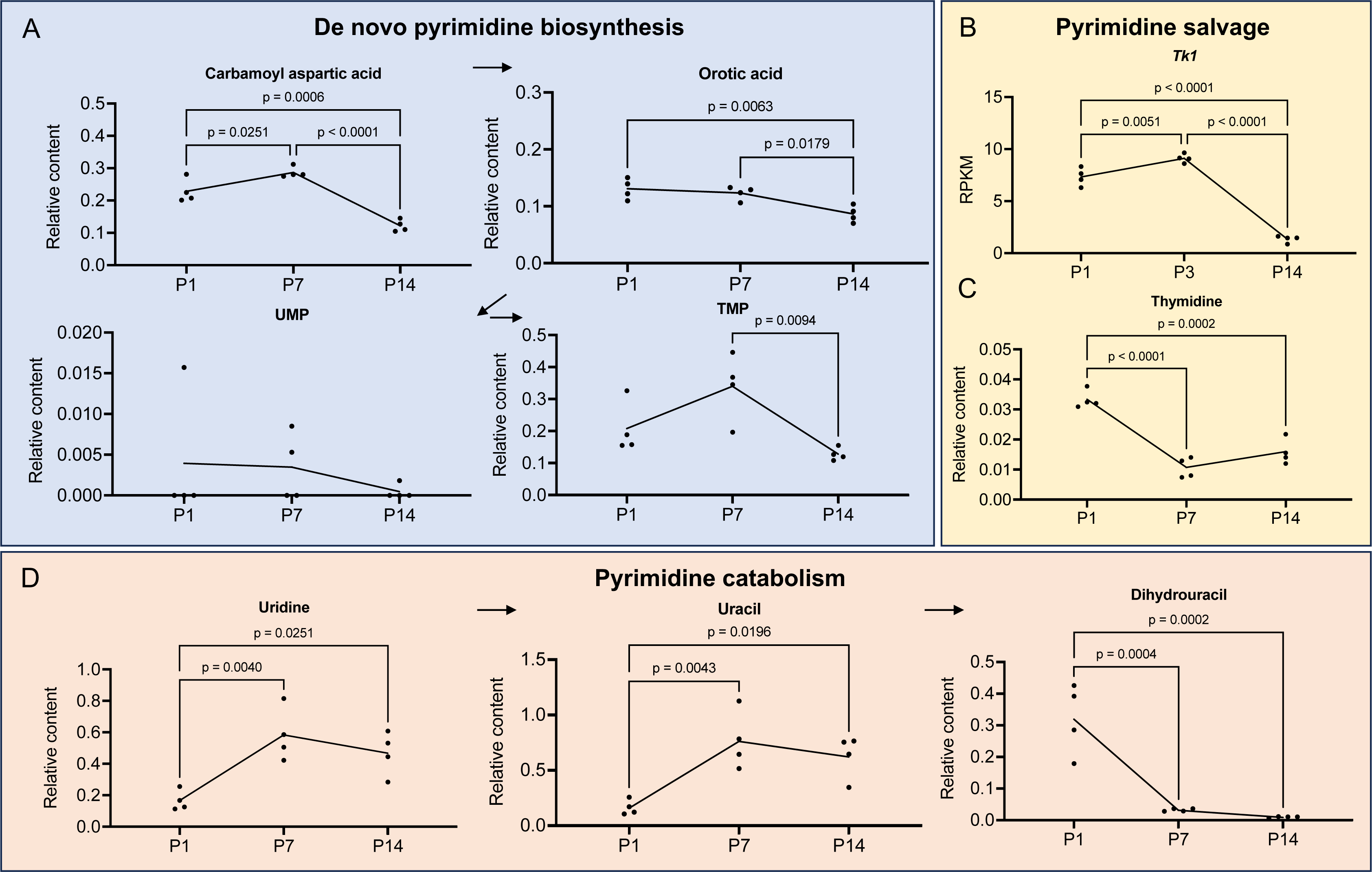
Changes of pyrimidine metabolism in the postnatal mouse heart. (A) Time-course analysis of metabolites related to the de novo pyrimidine biosynthesis in the mouse heart from P1 to P14. (B) Time-course analysis of expression of *Tk1*, a gene encoding thymidine kinase, in the heart from P1 to P14 (re-analyzed from Cardoso-Moreira, M. et al^56^). (C) Time-course analysis of thymidine, associated with the pyrimidine salvage pathway in the heart from P1 to P14. (D) Time-course analysis of metabolites related to the pyrimidine catabolism pathway in the mouse heart from P1 to P14.

**Supplementary Figure 6.**
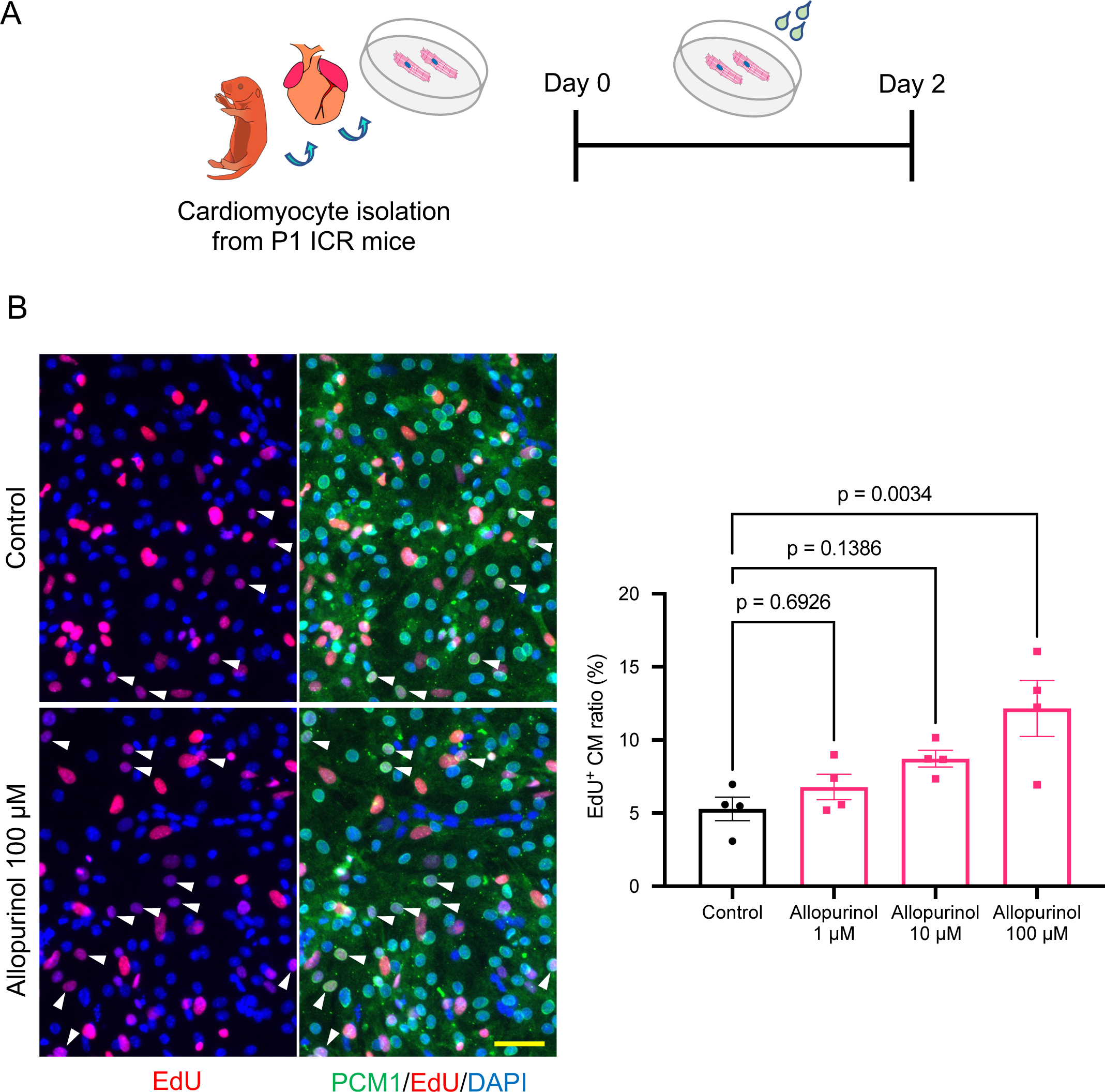
Effects of allopurinol treatment on neonatal cardiomyocyte proliferation. (A) A scheme depicting the protocol of NMVCs isolation from P1 mice and primary culture with allopurinol. (B) Representative images of EdU, PCM1 (cardiomyocyte nuclear marker), and DAPI (left), and DNA synthesis in cardiomyocytes evaluated by the rate of EdU^+^ cardiomyocyte nuclei (right). Quantification was conducted on 500 x 500 µm^2^ area across 4 wells in each group. In the representative images, white arrowheads indicate EdU^+^ cardiomyocyte nuclei. A scale bar indicates 50 µm.

**Supplementary Figure 7.**
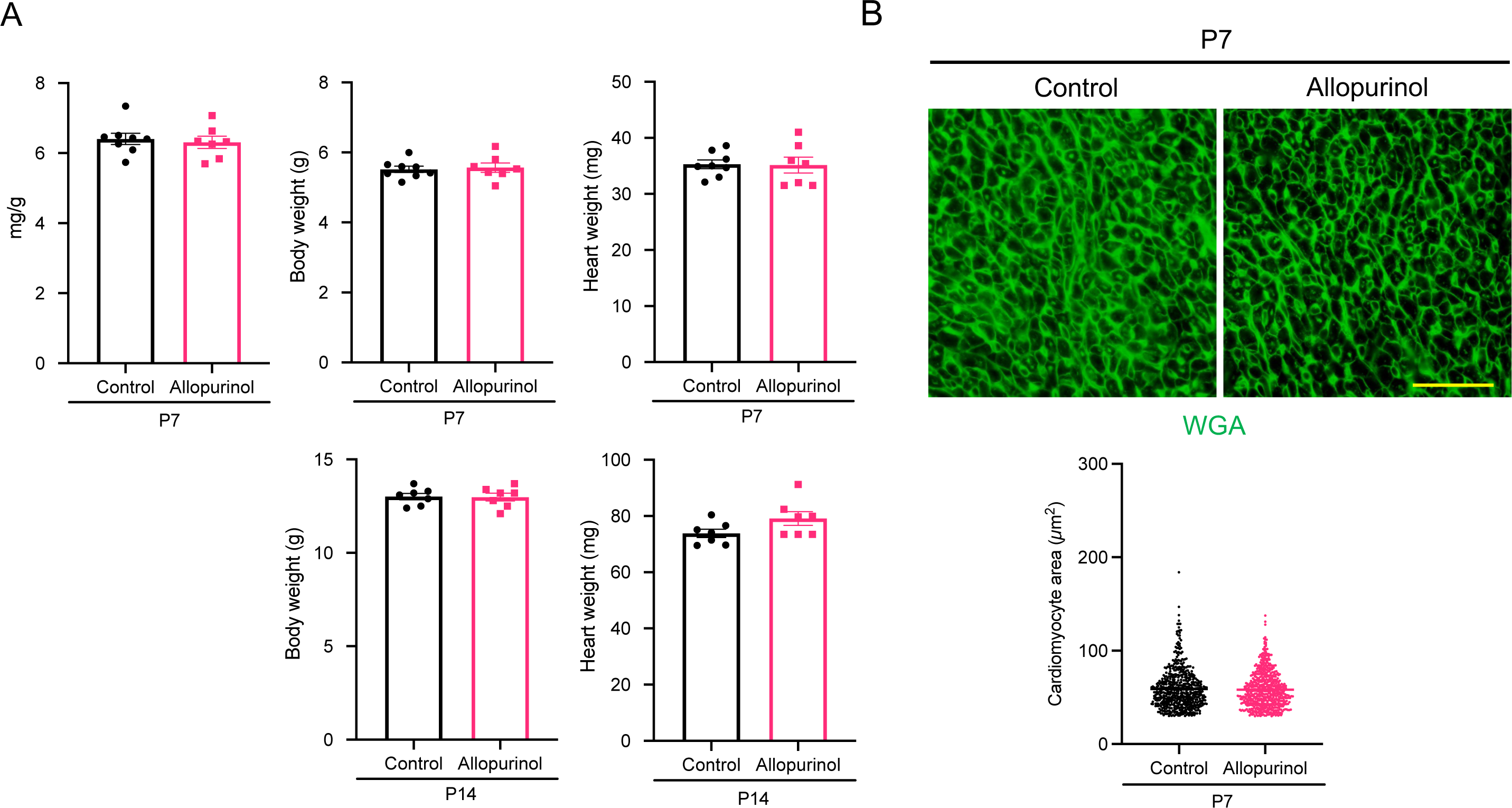
Effects of allopurinol treatment on the postnatal mouse heart. (A) The ratio of heart weight per body weight at P7, and body weight and heart weight at P7 and P14 in control and allopurinol-treated mice. (B) Cardiomyocyte cell size evaluated by the imaging of WGA stained sections of the heart from control and allopurinol-treated mice. A scale bar indicates 50 µm.

**Supplementary Table 1 Metabolome analysis of the postnatal mouse and opossum heart**

**Supplementary Table 2 Integrated omics analysis**

**Supplementary Table 3 Metabolome analysis of the heart from control and NAC-treated mice**

**Supplementary Table 4 Metabolome analysis of the heart from control and DCA-treated mice**

**Supplementary Table 5 Metabolome analysis of the freeze-clamped mouse heart**

